# Structural basis of endogenous inverse agonism and subtype selectivity at melanocortin receptors

**DOI:** 10.64898/2026.09.03.748715

**Authors:** Qinxin Sun, Shuhao Zhang, Xin Zhang, Xiaoou Sun, Xiangyu Liu

## Abstract

Endogenous inverse agonists suppress constitutive G protein-coupled receptor (GPCR) signaling, but their mechanisms remain poorly understood. Here we report cryo-EM structures of melanocortin-1 receptor (MC1R) bound to its endogenous inverse agonist, agouti signaling protein (ASIP) and melanocortin-4 receptor (MC4R) bound to agouti-related protein (AgRP). Together with previously reported structures and those we determined using extracellular nanobodies developed here, these data delineate a conformational continuum underlying receptor activation and silencing. Both inverse agonists occlude the orthosteric pocket as molecular corks and drive a shared TM3-centered extracellular remodeling. Against this common mechanism, subtype selectivity is encoded not by the conserved orthosteric pocket but by the divergent extracellular receptor surface, engaged through an ASIP C-terminal loop-dependent clasp. These findings establish a mechanistic framework for endogenous inverse agonism and identify the receptor periphery as a tractable target for subtype-selective modulation.

## Introduction

G protein-coupled receptors (GPCRs) sense extracellular signals and mediate cellular responses, thereby regulating physiological processes. While most endogenous GPCR ligands function as agonists, endogenous antagonists or inverse agonists have been identified in a few physiological systems, exemplified by the melanocortin-targeting agouti proteins^1,2^ and the ghrelin-receptor regulator liver-expressed antimicrobial peptide 2 (LEAP2)^3^. Among these, agouti signaling protein (ASIP)^4^ and agouti-related protein (AgRP)^5,6^, which target the melanocortin receptor (MCR) family, constitute the best-characterized endogenous inverse agonist system. However, the molecular mechanisms by which these endogenous inverse agonists regulate receptor function remain elusive.

The MCR family consists of five Gs-coupled subtypes (MC1R-MC5R) that regulate pigmentation, steroidogenesis, energy homeostasis, and exocrine gland function^7^. These receptors are activated by proopiomelanocortin (POMC)-derived peptides, including adrenocorticotropic hormone (ACTH) and the melanocyte-stimulating hormones (α-, β-, and γ-MSH), which share a conserved His-Phe-Arg-Trp (HFRW) pharmacophore that engages a highly conserved orthosteric pocket^7^. Several MCRs also exhibit appreciable constitutive activity, establishing a basal signaling tone that is suppressed by endogenous inverse agonists. Among these, MC1R displays notably high constitutive cAMP signaling activity^8^, potentially sustained by an N-terminal tethered agonist^9^. This basal signaling promotes eumelanin synthesis and UV photoprotection in melanocytes, and its disruption is associated with the red hair phenotype, fair and UV-sensitive skin, and increased melanoma susceptibility^10–12^. MC4R, expressed in hypothalamic neurons, is a principal regulator of appetite and energy expenditure^13,14^. Like MC1R, MC4R retains a constitutive activity that is suppressed by its endogenous inverse agonist^15^. Loss-of-function mutations in MC4R represent the most common monogenic cause of severe early-onset obesity^14^, and the MC4R agonist setmelanotide has been approved for the treatment of several genetic obesity disorders^16,17^.

ASIP and AgRP are secreted proteins composed of divergent N-terminal regions and a highly conserved C-terminal cysteine-rich domain that alone is sufficient for receptor inhibition^18–21^. This cysteine-rich domain adopts an inhibitor cystine knot (ICK) fold^22–25^, a compact disulfide-stabilized scaffold more commonly found in animal venom peptides such as cone snail conotoxins and spider venom toxins^26,27^. Each protein displays an Arg-Phe-Phe (RFF) motif on a loop projecting from the knot, which serves as its core receptor-binding pharmacophore^28–30^. The physicochemical resemblance between the RFF motif and the HFRW motif of melanocortin peptides, together with the mutually exclusive receptor binding, has long supported the model that agouti proteins function as simple competitive orthosteric antagonists^1,31,32^.

Despite these shared features, ASIP and AgRP diverge in their receptor preferences^1,2^. ASIP acts across several melanocortin subtypes with a preference for MC1R^1^, where it opposes α-MSH-mediated MC1R activation in the hair follicle to regulate the switch from eumelanin to pheomelanin production^32,33^. AgRP, by contrast, is expressed in hypothalamic neurons and potently inhibits MC3R and MC4R^2^ to promote feeding^34^, and only shows weak activity toward MC1R^2^. How agouti proteins sharing a conserved pharmacophore achieve such asymmetric receptor preferences has remained unknown.

Both MC1R and MC4R are attractive therapeutic targets, yet the high conservation of the orthosteric binding pocket across the MCR family has hindered the development of subtype-selective small-molecule and peptide ligands^35^. Nanobodies have emerged as promising molecular scaffolds to achieve superior selectivity among conserved GPCR subtypes, owing to their ability to engage larger and more divergent extracellular epitopes^36^. Selective nanobody modulators have been developed for several class A GPCRs, including the apelin receptor^37^, the angiotensin II type 1 receptor^36,38^, and the μ-opioid receptor^39^. Within the melanocortin receptor family, a functional nanobody agonist has been described for MC4R^40^. For MC1R, however, no selective nanobody inhibitor has been reported, despite its therapeutic relevance in pigmentary disorders and melanoma.

In this work, we combine cryo-electron microscopy (cryo-EM) analyses of the ASIP-MC1R and AgRP-MC4R complexes with structures of MC1R in multiple functional states to reveal the structural basis of endogenous inverse agonism and receptor selectivity in the melanocortin system. Both agouti proteins engage the orthosteric pocket through a conserved arginine-for-calcium substitution and induce a shared TM3-centered extracellular remodeling that stabilizes the inactive receptor. Yet against this shared mechanism, subtype selectivity is encoded not within the conserved orthosteric pocket but by an ASIP C-terminal loop-dependent clasp on the divergent ECL1/TM2 surface of MC1R. This explains why MC1R discriminates between the two ligands whereas MC4R does not. Consistent with this principle, we identify Nb96, a nanobody that independently converges on the same divergent ECL1 surface and acts as a subtype-selective antagonist of MC1R.

## Results

### Generation of a conformation-specific nanobody for ASIP-MC1R complex stabilization

Early attempts to determine the high-resolution structure of the ASIP-MC1R complex were limited by the conformational flexibility and the relatively small molecular size of the inactive-state receptor. We therefore sought a conformation-specific nanobody as a structural chaperone to stabilize the ASIP-bound inactive state and facilitate cryo-electron microscopy (cryo-EM) analysis.

Immune repertoires were generated by immunizing alpacas with an engineered inactive-state MC1R construct employing the Nb6 approach^41^, and the resulting immune repertoires were screened by yeast surface display system^42^. Following enrichment against inactive MC1R and subsequent selection for binding to the ASIP-bound receptor (Extended Data Fig. 1a), we identified Nb1, a high-affinity nanobody that selectively binds to and stabilizes the ASIP-MC1R complex (Extended Data Fig. 1b,c). Incorporation of Nb1 substantially improved particle homogeneity and enabled determination of the ASIP-MC1R-Nb6-Nb1 complex at 2.94 Å (Fig. 1a, Table 1 and Extended Data Fig. 2).

**Fig. 1.**
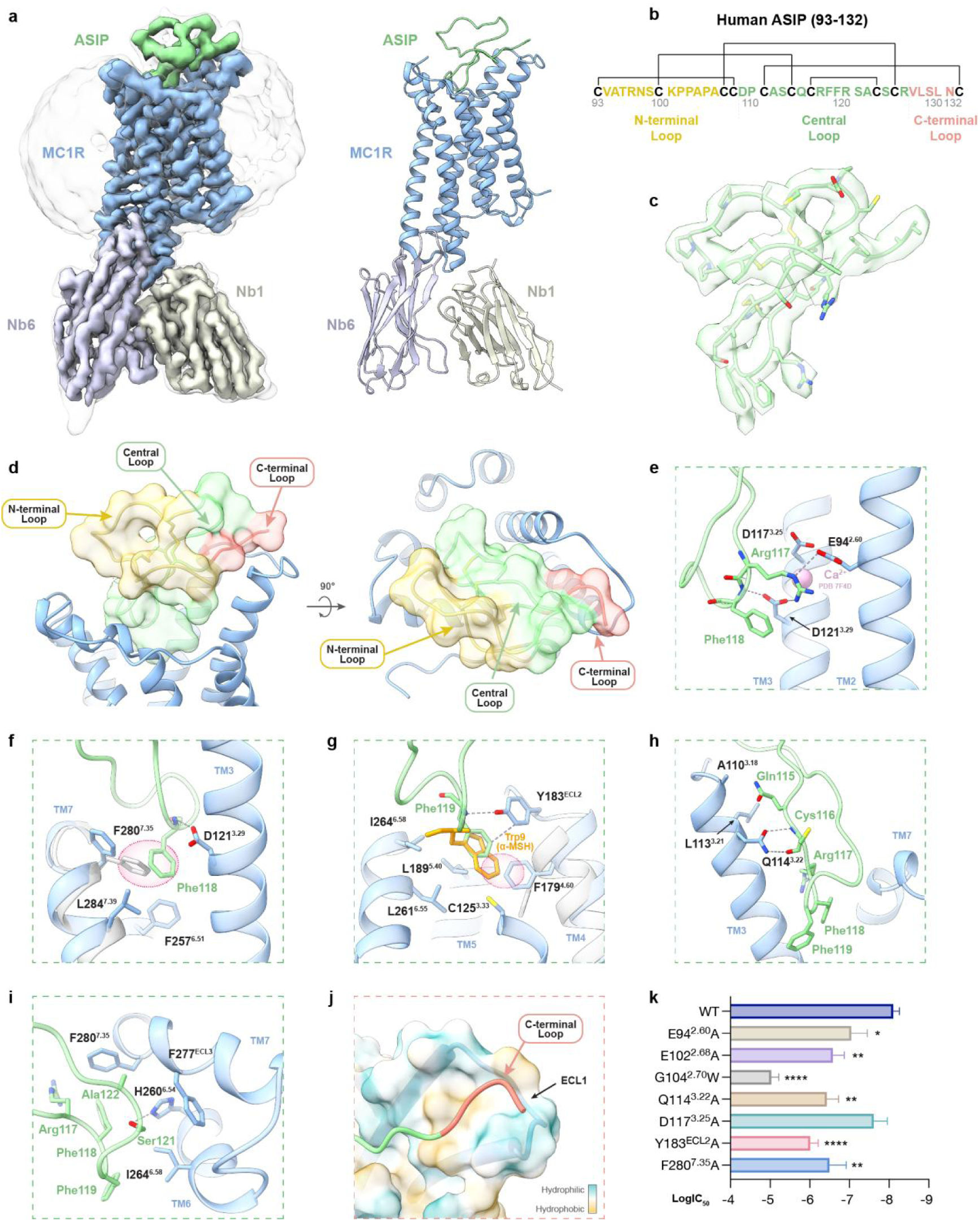
Cryo-EM structure of the ASIP-MC1R complex and recognition of the ASIP RFF motif. **a,** Cryo-EM density map (left) and model (right) of the ASIP-MC1R-Nb6-Nb1 complex. ASIP, green; MC1R, blue; Nb6, light purple; Nb1, pale green. The detergent micelle is shown as a transparent surface. **b,** Sequence of human ASIP (residues 93-132), with disulfide connectivity indicated above. The N-terminal, central and C-terminal loops are colored yellow, green and salmon, respectively. **c,** Cryo-EM density (mesh) for ASIP (residues 96-130). **d,** Two orthogonal views of the three ASIP loops (shown as surface; colored as in b) positioned over the extracellular vestibule of MC1R. **e-i,** Detailed interactions of the ASIP central loop within the MC1R orthosteric pocket. Arg117 engages the acidic triad E94^2.60^, D117^3.25^ and D121^3.29^. The Ca^2+^ position in α-MSH-bound MC1R (PDB 7F4D) is shown (pink sphere) **(e)**. Phe118 within the TM3/TM6/TM7 sub-pocket. F280^7.35^ in α-MSH-bound MC1R (gray) would sterically clash with Phe118 **(f)**. Phe119 at the TM4/TM5/ECL2 interface, superimposed with Trp9 (orange) of α-MSH, which would sterically clash with F179^4.60^ in the ASIP-bound state **(g)**. Interactions involving the Gln115-Cys116 N-terminal flank **(h)**. Interactions involving the Ser121-Ala122 C-terminal flank **(i)**. Interacting residues are shown as sticks and gray dashed lines indicate polar interactions. **j,** Close-up view of the ASIP C-terminal loop interacting with ECL1 of MC1R. The receptor surface is colored by hydrophobicity. Green, hydrophilic; yellow, hydrophobic. **k,** Effect of the indicated MC1R mutations on ASIP inhibitory potency (LogIC_50_), measured by GloSensor cAMP assay. Data are shown as mean ± SEM from 6 biologically independent experiments for WT MC1R and 3 biologically independent experiments for each mutant. Statistical significance of pIC_50_ was assessed by one-way ANOVA followed by Dunnett’s multiple-comparisons test versus WT MC1R. *P < 0.05, **P < 0.01, ***P < 0.001, ****P < 0.0001.

### Overall cryo-EM structure of the ASIP-MC1R complex

To maximize the chances of observing ASIP, we fused the ASIP fragment (residues 80–132) to the N-terminus of MC1R and also added separately purified ASIP protein (residues 23–132) (Extended Data Fig. 1d,e) to the system. Through these efforts, we obtained interpretable cryo-EM density for the C-terminal cysteine-rich domain (residues 96–130) (Fig. 1b,c). This observation is consistent with previous reports that this domain constitutes the minimal functional module sufficient for receptor binding^19,22^. In contrast, the N-terminal pro-region, which is characterized by intrinsic flexibility and considered dispensable for direct receptor engagement, remained unresolved, likely due to its high degree of conformational heterogeneity^22^.

The resolved ASIP fragment exhibits the characteristic ICK fold (Fig. 1c). Extending from its cystine-knot core are three polypeptide loops, the N-terminal loop (93-108), the central loop (109-126), and the C-terminal loop^22^ (127-132), which project outward to interact with the receptor (Fig. 1b,d). Superposition onto the free-state NMR ensemble (PDB 1Y7K) showed that the cystine-knot core of ASIP is retained upon binding, with differences confined to the peripheral loops, indicating that the cysteine-rich domain is pre-folded for MC1R engagement (Extended Data Fig. 3a).

Unlike melanocortin peptide agonists, which adopt a characteristic U-shaped conformation at the extracellular end of the transmembrane domain (TMD) and insert the conserved HxRW motif deep into the binding pocket (α-MSH, PDB 7F4D; afamelanotide, PDB 7F4H; SHU9119, PDB 7F4I)^35^, the cysteine-rich domain of ASIP acts as a molecular “cork”, positioning itself over the extracellular vestibule of MC1R to achieve comprehensive occlusion (Fig. 1d). In the central loop, two strands of an antiparallel β-sheet align parallel to TM3, anchoring the scaffold and functioning as a “plug” that extends downward into the TMD core. Positioned above this central anchor, the N-terminal loop sits at the outer edge, situated distal to the TMD core, and serves as a structural “lid” that extends occlusion to TM1/TM7. Meanwhile, the C-terminal loop extends laterally as a “clasp” against TM2 and ECL1 (Fig. 1d). Through this three-pronged architecture, ASIP spans the entire extracellular vestibule of MC1R rather than concentrating its contacts within the orthosteric core. The ASIP cysteine-rich domain is substantially larger than α-MSH, yet buries a smaller ligand-side interface (≈ 1020 Å^2^ for ASIP and ≈ 1155 Å^2^ for α-MSH, calculated with PDBePISA^43^). Only about one-third of the accessible surface of ASIP is buried, compared with approximately two-thirds for α-MSH, indicating that ASIP achieves receptor occlusion through broad but shallow surface engagement.

### Recognition of the RFF pharmacophore by the MC1R orthosteric pocket

The central loop harbors the conserved RFF tripeptide, which is well-established as the core pharmacophore of ASIP^30^. Our structure reveals that this RFF motif penetrates deeply into the MC1R orthosteric pocket, engaging residues within the TM2–TM4 region (Fig. 1d). Compared with the HFRW pharmacophore of α-MSH, the RFF motif exhibits a physicochemical resemblance, with both motifs sharing a comparable arrangement consisting of a positively charged residue and an aromatic cluster. However, structural superposition showed that the two motifs occupy distinct regions of the binding pocket (Extended Data Fig. 3b). Consistent with this observation, residue-level interaction analysis demonstrated largely non-overlapping receptor contact networks for the two pharmacophores (Extended Data Fig. 3c,d), implying that the RFF motif interacts with MC1R through a binding mode distinct from that of α-MSH.

The guanidinium group of Arg117 projects into the interhelical cleft between TM2 and TM3, where it forms a salt-bridge network with the acidic triad E94^2.60^, D117^3.25^ and D121^3.29^ (Fig. 1e). Alanine substitution of E94^2.60^ markedly diminishes ASIP-mediated inhibition (Fig. 1k, Extended Data Fig. 3h and Extended Data Fig. 4). These acidic residues are highly conserved across the melanocortin receptor family and constitute a Ca^2+^ coordination site within the orthosteric peptide-binding pocket^35,44^. In previously reported agonist-bound active states of MC1R and related melanocortin receptors, well-resolved calcium ion density is observed at this site, coordinated by the acidic residues and together with backbone atoms of the melanocortin peptide, consistent with the established role of Ca^2+^ as an essential cofactor for agonist recognition and receptor activation^45^. Interestingly, no calcium density is observed at this site in the ASIP-bound structure. Instead, the elongated and positively charged side chain of Arg117 directly engages the acidic triad, functionally substituting for the Ca^2+^ and compensating for the local cationic requirement within the binding pocket (Fig. 1e).

Analogous to Phe7 of the α-MSH pharmacophore, Phe118 resides at the base of the orthosteric binding pocket, yet engages this region through a markedly distinct geometry (Fig. 1f). The insertion depth of Phe118 is considerably shallower than that of Phe7 (Extended Data Fig. 3e). Furthermore, rather than orienting toward TM3 like Phe7, the benzyl side chain of Phe118 projects directly into the interhelical cleft formed by TM3, TM6, and TM7. Within this sub-pocket, Phe118 establishes extensive hydrophobic and van der Waals interactions with F257^6.51^, F280^7.35^ and L284^7.39^ (Fig. 1f). In addition, the main-chain amide of Phe118 forms a hydrogen bond with D121^3.29^, providing a polar backbone anchor that complements the Arg117-acidic triad network. Notably, the proximity of Phe118 to TM7 forces F280^7.35^ to undergo a pronounced outward displacement in the ASIP-bound complex. In contrast, in the α-MSH-bound structure, F280^7.35^ adopts an inward-facing conformation, which would otherwise result in a steric clash with Phe118 (Fig. 1f).

Adjacent to Phe118, Phe119 further anchors the RFF motif within the orthosteric pocket, near TM4, TM5 and ECL2 (Fig. 1g). Its backbone amide forms a hydrogen bond with Y183^ECL2^, complemented by an edge-to-face π–π contact, consistent with the markedly reduced ASIP inhibitory potency of the Y183^ECL2^A mutant (Fig. 1k, Extended Data Fig. 3h and Extended Data Fig. 4). Phe119 further establishes hydrophobic contacts with F179^4.60^, L189^5.40^, L261^6.55^ and I264^6.58^, together with a sulfur–π interaction with C125^3.33^ (Fig. 1g). Despite occupying a similar position to the pyrrole moiety of Trp9 in α-MSH, Phe119 inserts more shallowly due to the smaller steric footprint of its benzyl ring relative to the bicyclic indole. This allows greater conformational flexibility for surrounding receptor residues. Consequently, F179^4.60^ is oriented toward Phe119 in the ASIP-bound structure, but adopts an outward conformation in the α-MSH-bound state to accommodate the bulkier indole group of Trp9 (Fig. 1g).

Flanking the deeply inserted RFF motif, the central loop extends across the extracellular receptor surface and establishes additional contacts with TM3, TM6, TM7, and ECL3. On the N-terminal flank, Cys116 and Gln115 both engage Q114^3.22^ at the extracellular end of TM3. The backbone of Cys116 forms bidentate hydrogen bonds with Q114^3.22^, reinforcing the β-strand preceding the RFF motif and helping position Arg117 (Fig. 1h). Consistent with these interactions, alanine substitution at Q114^3.22^ reduced ASIP inhibitory potency by approximately 50-fold (Fig. 1k, Extended Data Fig. 3h and Extended Data Fig. 4). On the C-terminal flank, Ser121 engages a polar and hydrophobic cluster at the extracellular end of TM6 and ECL3 (Fig. 1i). It forms a hydrogen bond with H260^6.54^ while also packing against I264^6.58^ and F277^ECL3^. Adjacent Ala122 further contributes to hydrophobic packing with F277^ECL3^ and F280^7.35^ (Fig. 1i). Mutation of F280^7.35 likewise reduced ASIP inhibitory potency, supporting the contribution of this peripheral interaction network (Fig. 1k, Extended Data Fig. 3h and Extended Data Fig. 4).

### Extracellular occlusion of MC1R by three structurally distinct ASIP loops

Whereas the central loop achieves receptor recognition through deep penetration and an extensive contact network, the N-terminal loop (residues 93-108, with 96-108 resolved) establishes only negligible buried contacts (≈ 15 Å^2^) and remains suspended above the orthosteric vestibule (Fig. 1d). Rather than redundantly covering the central loop region, the N-terminal “lid” extends the occlusion into the TM1/TM7 sector that remains broadly exposed in the α-MSH-bound state (Extended Data Fig. 3f).

The C-terminal loop (residues 127-132, with 127-130 resolved) extends laterally from the cysteine-rich scaffold to engage the extracellular end of TM2 together with ECL1 of MC1R, completing the three-pronged engagement of ASIP (Fig. 1d). While the central loop “plug” buries approximately 811 Å^2^ through deep insertion and the N-terminal “lid” hovers over the vestibule with negligible buried contacts, the C-terminal loop establishes an intermediate interface of ∼194 Å^2^ (Fig. 1d). Despite this limited footprint, it contributes to interactions previously identified as critical for ASIP binding^28^. This interface is dominated by hydrophobic contacts (Fig. 1d and Extended Data Fig. 3g), and consistent with its close-packed nature, substitution of G104^ECL1^ with a bulky tryptophan reduces ASIP affinity to the micromolar range (Fig. 1k, Extended Data Fig. 3h and Extended Data Fig. 4).

Together, the three loops of the ASIP cysteine-rich domain form a spatially complementary receptor engagement architecture, with each segment contributing through a distinct mode of interaction. This multilayered organization establishes ASIP as an extracellular occlusion module of MC1R, engaging the receptor through a fundamentally different binding strategy from that of peptide agonists.

### Structural basis of ASIP-mediated inverse agonism at MC1R

Among the five melanocortin receptor subtypes, MC1R exhibits the highest basal cAMP activity, potentially sustained by an N-terminal tethered agonist proposed to occupy the orthosteric pocket^8,9^. This intrinsic activity raises a central mechanistic question regarding ASIP-mediated inverse agonism: how does ASIP displace the tethered activator and suppress the conformational transitions underlying constitutive signaling? To define the structural basis of this process, we determined the Gs-uncoupled, ligand-free structure of MC1R stabilized by Nb18, an extracellular nanobody identified in this study, at 3.42 Å resolution (Extended Data Fig. 5, Extended Data Fig. 6 and Table 1). Nb18 is functionally silent, neither activating nor antagonizing α-MSH-stimulated cAMP responses (Extended Data Fig. 5b), and therefore captures MC1R in a resting-state configuration without G protein-induced stabilization. Together with the α-MSH-bound MC1R-Gs complex, the unliganded MC1R-Gs complex, and the ASIP-MC1R complex described here, these four structures define a conformational landscape spanning the resting, constitutively active, agonist-bound active, and inverse-agonist-stabilized states of MC1R.

In the agonist-bound state, α-MSH adopts a U-shaped conformation in the orthosteric cavity (Fig. 2a, left). By contrast, in the unliganded state, the receptor N-terminus likely occupies this region to sustain constitutive signaling (Fig. 2a, right). When ASIP binds, its cysteine-rich domain fills the binding pocket and blocks access for agonists (Fig. 2a). Consistent with this, in the ASIP-bound complex, the receptor N-terminus is displaced from the pocket and adopts a peripheral trajectory along TM7, similar to its position in the α-MSH-bound structure (Fig. 2b,c). Thus, ASIP acts as a steric blocker, preventing both the tethered agonist and α-MSH from binding.

**Fig. 2.**
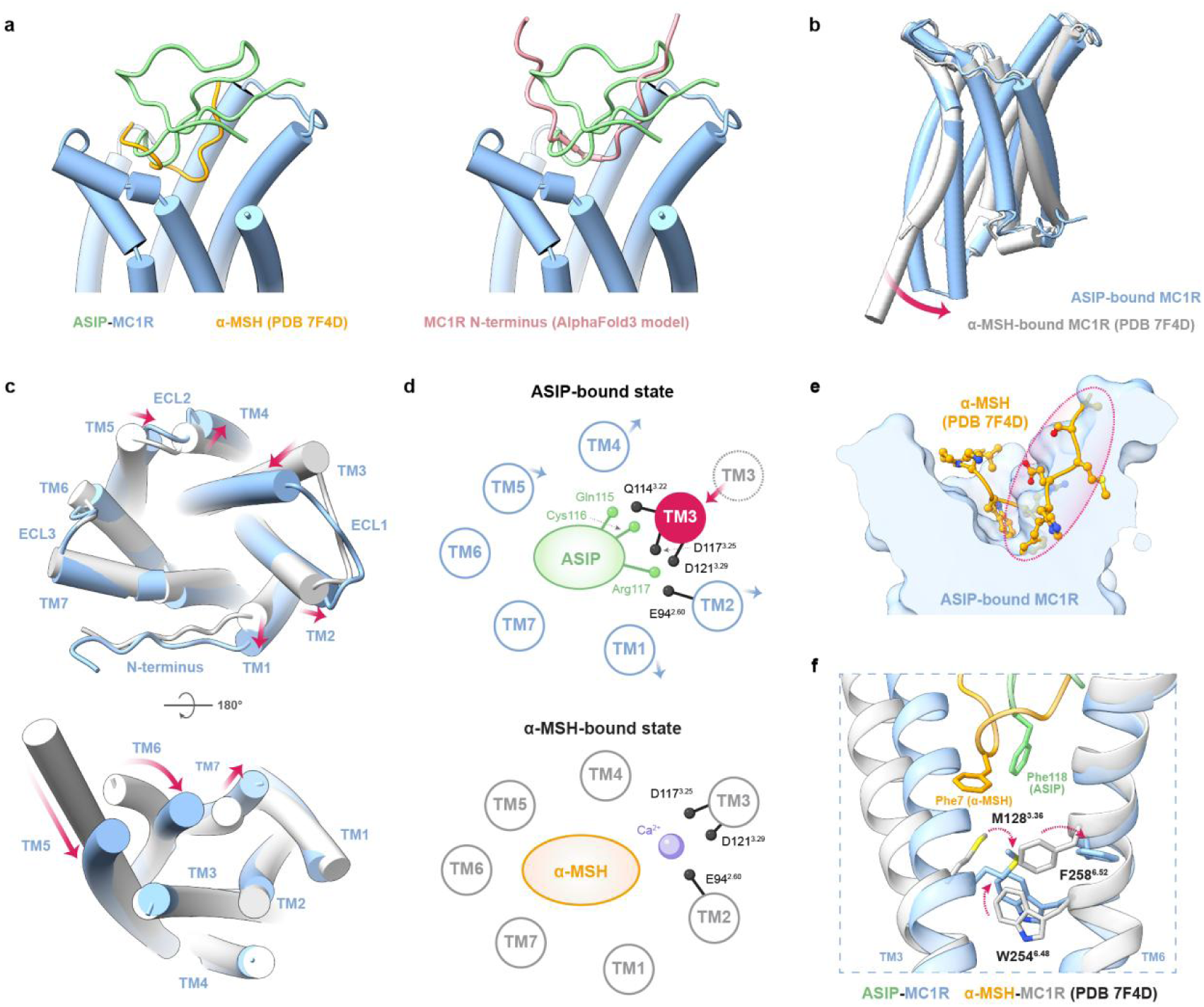
Structural basis of ASIP-mediated inverse agonism at MC1R. **a,** Binding positions within the MC1R orthosteric pocket. Left, ASIP (green) and α-MSH (orange; PDB 7F4D); right, the MC1R N-terminus (residues 1-32; salmon; AlphaFold3 model) are shown relative to the ASIP-bound receptor (blue). **b,** Superposition of ASIP-bound MC1R and α-MSH-bound MC1R (gray); arrows indicate inward displacement of TM6. **c,** Extracellular (top) and intracellular (bottom; 180° rotation) views of transmembrane helix rearrangements between ASIP-bound and α-MSH-bound MC1R. Arrows indicate the direction of displacement. **d,** ASIP-induced inward movement of TM3. In the ASIP-bound state (top), Arg117, together with the Gln115-Cys116 segment, anchors the extracellular end of TM3. In the α-MSH-bound state (bottom), this interaction network is replaced by Ca^2+^-mediated coordination involving α-MSH and the acidic triad. **e,** α-MSH superimposed onto the surface of the ASIP-bound MC1R orthosteric pocket, illustrating steric incompatibility with the remodeled pocket. **f,** Conformations of the toggle switch (M128^3.36^, W254^6.48^ and F258^6.52^) in ASIP-bound and α-MSH-bound MC1R. Phe7 of α-MSH and Phe118 of ASIP are shown. Residues are shown as sticks.

Superposition of the ASIP-MC1R complex with the α-MSH-bound MC1R-Gs structure reveals global extracellular and intracellular remodeling (Fig. 2b,c). Intracellularly, the cytoplasmic end of TM6 moves inward relative to the active state, consistent with the conserved features of inactive class A GPCRs (Fig. 2b,c). Extracellularly, TM1-TM4 undergo distinct rearrangements, with the most prominent and unusual change occurring at TM3 (Fig. 2c). Its extracellular end shifts inward by 3.8 Å at V112^3.20^ relative to the agonist-bound state, exceeding the shifts seen in the constitutively active state (1.2 Å) and the Nb18-stabilized ligand-free state (2.5 Å) (Extended Data Fig. 7a). This progressive inward movement across the agonist-bound, constitutively active, resting, and ASIP-bound states defines a continuum of extracellular TM3 remodeling, with ASIP binding driving the receptor beyond the resting-state configuration and into the inverse-agonist-stabilized state. Such a pronounced inward movement is atypical of class A GPCRs^46^, as representative receptors including theβ_2_-adrenergic receptor and the μ-opioid receptor show no comparable inward displacement between active and inactive states (Extended Data Fig. 7b,c). Therefore, the unusual inward displacement of TM3 observed here appears to underlie the broader extracellular reshaping in ASIP-bound MC1R.

The inward displacement of TM3 is stabilized by a specific multi-anchor clamp formed by ASIP. As described above, Arg117 of the RFF motif engages the conserved acidic triad E94^2.60^, D117^3.25^ and D121^3.29^ through a salt-bridge network, while the Gln115-Cys116 segment further anchors ASIP to TM3 through polar contacts (Fig. 1e,h and Fig. 2d, top). In contrast, α-MSH uses a coordinated Ca^2+^ ion to mediate interactions with the same TM3 acidic cluster (Fig. 2d, bottom), a feature retained even in the ligand-free resting state (Extended Data Fig. 7d). By replacing this spherical ion with a directional three-residue clamp, ASIP converts a metal-coordination center into a tethered interaction network (Fig. 2d). This structural change explains why TM3 shifts further inward than in the Ca^2+^-coordinated resting state.

Although TM1, TM2, and TM4 move outward, the inward displacement of TM3 narrows the orthosteric pocket. This new shape prevents the binding of both melanocortin agonists and the Ca^2+^ cofactor. Overlays show that the inward-shifted TM3 backbone directly clashes with both the N-terminal segment of α-MSH (before Arg8) and the Ca^2+^ ion (Fig. 2e). Additionally, side chains of F280^7.35^ and F179^4.60^ adopt rotamers in the ASIP-bound pocket that are incompatible with α-MSH (Fig. 1f,g). Therefore, ASIP acts as an inverse agonist not just by blocking the orthosteric pocket, but also by stabilizing MC1R in a conformation that physically excludes the endogenous activators.

Beyond the extracellular remodeling, the ASIP-bound receptor is captured in an inactive intracellular conformation. When α-MSH binds, its Phe7 from the HFRW pharmacophore stabilizes the toggle-gating residue M128^3.36^ in an upward rotamer, which in turn induces a downward displacement of the toggle switch residue W254^6.48^, triggering the canonical activation cascade by rearranging conserved microswitch motifs (Fig. 2f). In contrast, ASIP’s RFF motif inserts less deeply and does not directly engage M128^3.36^ or W254^6.48^. Instead, M128^3.36^ adopts a downward rotamer (Fig. 2f), which forces F258^6.52^ to rotate outward to avoid a clash. The same downward arrangement of M128^3.36^ is observed in the ligand-free resting state of MC1R, whereas the previously reported constitutively active MC1R structure displays an intermediate conformation (Extended Data Fig. 7e). Furthermore, the inward shift of extracellular TM3 is propagated along the helix, translating the M128^3.36^ Cα downward by 2.6 Å relative to the α-MSH-bound state (Fig. 2f). Thus, the entire 3.36-6.48 interface is repositioned by a backbone-level displacement rather than by a side-chain rotameric change. Consequently, W254^6.48^ is held in its inactive upward orientation despite the absence of direct ligand contact with the toggle switch residues. Downstream, the receptor displays the hallmarks of an inactive class A GPCR. In the MLF motif (M199^5.50^, L132^3.40^, F250^6.44^), F250^6.44^ points upward, and the DRY motif retains the D141^3.49^-R142^3.50^ salt bridge with Y143^3.51^ oriented away from TM5 (Extended Data Fig. 7f).

Collectively, these findings explain how ASIP acts as an inverse agonist at MC1R through a multilayered mechanism. ASIP physically blocks the orthosteric pocket, keeping out both the receptor’s N-terminal tethered agonist and endogenous melanocortin peptides. In parallel, its cysteine-rich domain reshapes the extracellular part of MC1R into a constricted and agonist-incompatible conformation driven by the inward displacement of TM3. This helical rearrangement, together with the shallow insertion of the RFF motif, relays the extracellular binding event to the toggle switch, which is held in its inactive conformation without direct ligand contact, so that the conformational transition required for activation is not initiated. Together, these features stabilize MC1R in an inactive conformation that is refractory both to activation by endogenous agonists and to intrinsic constitutive activity.

### Conserved and divergent structural features of AgRP recognition by MC4R

Originally identified to stabilize the ASIP-bound MC1R complex, Nb1 recognizes the conserved intracellular surface formed by Nb6 and the chimeric κ-opioid receptor (KOR) ICL3, enabling it to serve as a transferable intracellular chaperone for the structure determination of Nb6-based inactive-state GPCRs (Extended Data Fig. 8a). Leveraging this transferability, we determined the AgRP-MC4R complex by cryo-EM at a global resolution of 3.01 Å (Fig. 3a, Extended Data Fig. 8b-h and Table 1). The resolved AgRP fragment (residues 89-122) encompasses the C-terminal cysteine-rich domain (Fig. 3c), consistent with previous reports that this region constitutes the minimal receptor-binding module^21,24^. This domain adopts an inhibitor cystine knot fold that closely resembles the solution NMR structure (Extended Data Fig. 9a). As observed for ASIP bound to MC1R, it functions as a molecular cork positioned over the extracellular vestibule of MC4R (Fig. 3d). Extending from the cystine knot, the N-terminal loop (residues 87-104) and the central loop (residues 105-120) engage the receptor, whereas the C-terminal loop (residues 121-132) was largely unresolved, with only residues 121-122 modeled (Fig. 3d,j).

**Fig. 3.**
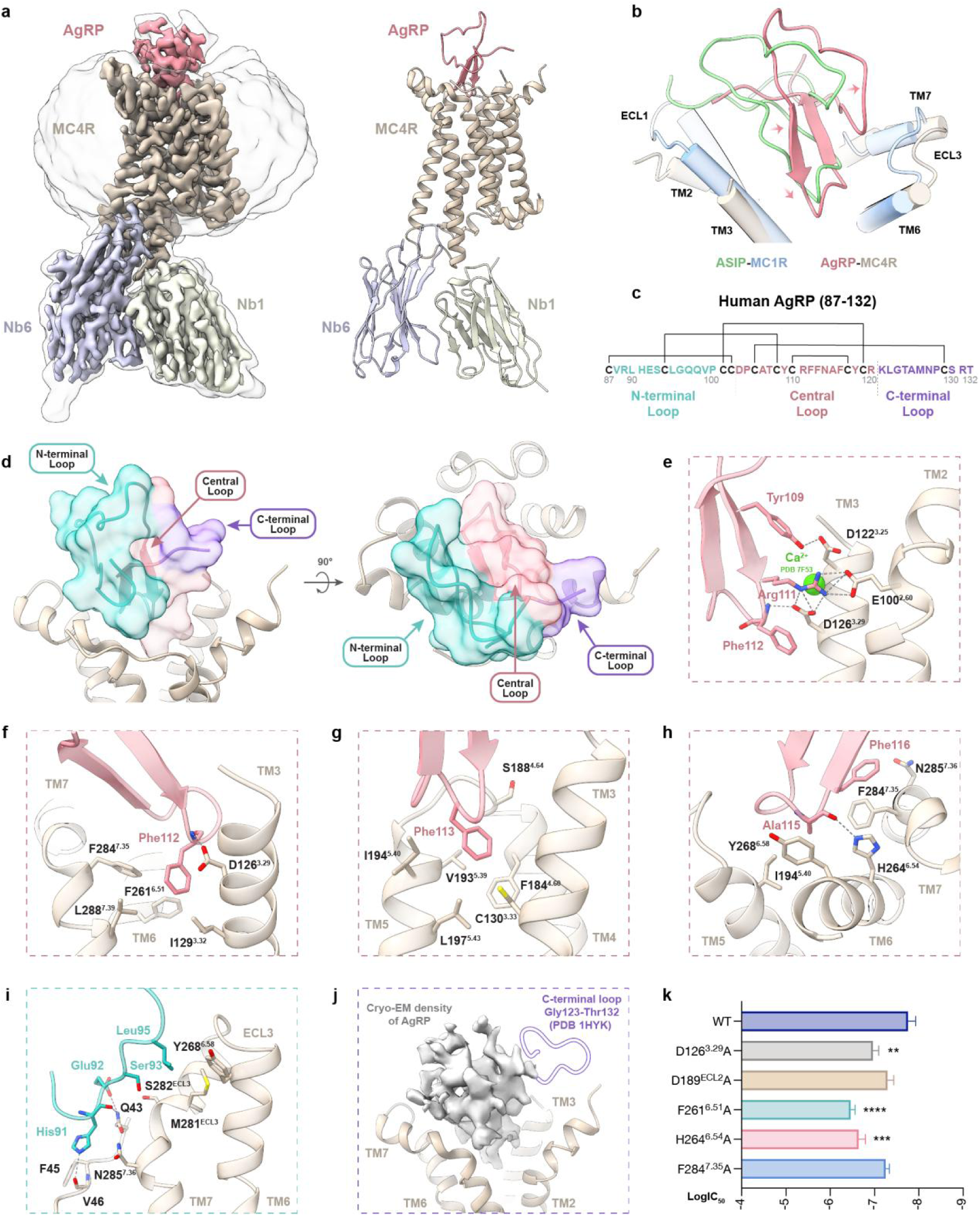
Conserved pharmacophore recognition and divergent loop engagement in the AgRP-MC4R complex. **a,** Cryo-EM density map (left) and model (right) of the AgRP-MC4R-Nb6-Nb1 complex. AgRP, red; MC4R, tan; Nb6, light purple; Nb1, pale green. The detergent micelle is shown as a transparent surface. **b,** Superposition of the AgRP-MC4R and ASIP-MC1R complexes, showing the relative position of the two cysteine-rich domains within the extracellular vestibule. Arrows indicate the displacement of AgRP relative to ASIP. **c,** Sequence and disulfide connectivity of human AgRP (residues 87–132). The N-terminal, central and C-terminal loops are colored cyan, red and purple, respectively. Black lines denote disulfide bonds. **d,** Surface representation of the three AgRP loops positioned over the extracellular vestibule of MC4R, shown in two orthogonal views. Colors as in (**c**). **e-h,** Detailed interactions of the AgRP central loop within the MC4R orthosteric pocket. Tyr109 and Arg111 engage the acidic triad E100^2.60^, D122^3.25^ and D126^3.29^ of MC4R. The Ca^2+^ position in α-MSH-bound MC4R (PDB 7F53) is shown (green sphere) (**e**). Phe112 within the deep aromatic sub-pocket formed by TM3, TM6 and TM7 (**f**). Phe113 within the shallow sub-pocket formed by TM4, TM5 and ECL2 (**g**). Interactions of Ala115 and Phe116 with the extracellular tips of TM6, TM7 and ECL3 (**h**). Interacting residues are shown as sticks and gray dashed lines indicate polar interactions. **i,** Interactions of the AgRP N-terminal loop with the MC4R N-terminus and the extracellular tips of TM6 and TM7. Interacting residues are shown as sticks and gray dashed lines indicate polar interactions. **j,** The AgRP C-terminal loop beyond Leu122 is not resolved in the cryo-EM density and was not modeled. **k,** LogIC_50_ values of AgRP-mediated inhibition of α-MSH-stimulated signaling at wild-type MC4R and the indicated mutants, measured by GloSensor cAMP assay. Data are shown as mean ± SEM from 5 biologically independent experiments for WT MC4R and 3 biologically independent experiments for each mutant. Statistical significance of pIC_50_ was assessed by one-way ANOVA followed by Dunnett’s multiple-comparisons test versus WT MC4R. *P < 0.05, **P < 0.01, ***P < 0.001, ****P < 0.0001.

As in ASIP, the AgRP central loop contains two strands of an antiparallel β-sheet and inserts into the orthosteric core as a plug (Fig. 3d). However, it adopts a more vertical orientation, tilted by ∼30° relative to ASIP, with its axis running more steeply into the transmembrane core and its peripheral end leaning toward the TM5–TM6 face (Fig. 3b; Extended Data Fig. 9b). The central loop harbors the Arg111-Phe112-Phe113 motif, which is functionally equivalent to the ASIP R117-F118-F119 pharmacophore and closely recapitulates its overall recognition mode (Extended Data Fig. 9c,d).

Similar to the ASIP-MC1R complex, no Ca^2+^ density is observed in the AgRP-MC4R structure. Instead, the conserved acidic triad E100^2.60^, D122^3.25^ and D126^3.29^ is engaged directly by Arg111 from AgRP (Fig. 3e). Consistent with this structural arrangement, alanine substitution of D126^3.29^ markedly reduced the inhibitory potency of AgRP (Fig. 3k and Extended Data Fig. 10a,b), whereas extracellular Ca^2+^, which potentiates α-MSH, does not affect AgRP binding^44^. However, the mode of triad engagement differs between the two complexes. Arg117^ASIP^ engages all three acidic residues of the MC1R triad (Fig. 1e), whereas Arg111^AgRP^ forms salt bridges with E100^2.60^ and D126^3.29^, while Tyr109^AgRP^ contributes an additional hydrogen bond to D122^3.25^ (Fig. 3e). Despite these distinct recognition modes, the underlying arginine-for-calcium interaction is conserved in both complexes, revealing a chemical principle shared by endogenous melanocortin inverse agonists.

Phe112^AgRP^ and Phe113^AgRP^ occupy the dual aromatic sub-pockets analogous to those engaged by Phe118^ASIP^ and Phe119^ASIP^, respectively. Phe112^AgRP^ inserts into the conserved hydrophobic cage formed by I129^3.32^, F261^6.51^, F284^7.35^ and L288^7.39^, with its main-chain amide hydrogen bonding to D126^3.29^ and providing a polar backbone anchor that complements the Arg111^AgRP^ interaction network (Fig. 3f). Although Phe112^AgRP^ penetrates modestly deeper than Phe118^ASIP^ (Extended Data Fig. 9c), its insertion remains substantially shallower than that of Phe7 in α-MSH (Extended Data Fig. 9e). Likewise, Phe113^AgRP^ extends deeper than its ASIP counterpart, Phe119^ASIP^, approaching the depth of Phe112^AgRP^ (Extended Data Fig. 9c). Nevertheless, it remains positioned within the TM4-TM5-ECL2 sub-pocket, where its aromatic ring forms a sulfur–π contact with Cys130^3.33^ (Fig. 3g), echoing the Cys^3.33^ sulfur–π interaction observed at MC1R (Fig. 1g).

Beyond the RFF core, the C-terminal flank of the central loop establishes additional contacts with TM6 and TM7 (Fig. 3h). In particular, the main-chain carbonyl of Ala115^AgRP^ hydrogen bonds with H264^6.54^, mirroring the Ser121^ASIP^-His260^6.54^ interaction observed in the ASIP-MC1R complex, while the adjacent Phe116^AgRP^ packs against Phe284^7.35^ at the extracellular tip of TM7. Consistent with the structural analysis, alanine substitution of the conserved anchor H264^6.54^ markedly reduced AgRP inhibitory potency (Fig. 3k and Extended Data Fig. 10a,b). On the ligand side, simultaneous alanine substitution of the Arg111-Phe112-Phe113 motif greatly reduced AgRP inhibitory activity, confirming the central loop as its functional core (Extended Data Fig. 10c, left).

In contrast to the conserved binding mode of the central loops, the N-terminal loops of AgRP and ASIP adopt markedly different architectures. Whereas the ASIP N-terminal loop lies above the MC1R extracellular vestibule without receptor contacts (Fig. 1d), the AgRP N-terminal loop forms an ordered contact network with the MC4R N-terminus and the extracellular tips of TM6 and TM7, where His91, Glu92, Ser93 and Leu95 form polar and hydrophobic contacts (Fig. 3i). Consistent with a functional contribution from this engagement, simultaneous alanine substitution of these four residues markedly reduced AgRP-mediated inhibition (Extended Data Fig. 10c, right). In contrast to this well-defined N-terminal engagement, the AgRP C-terminal loop remains largely unresolved beyond Leu122 (Fig. 3d,j), indicating that this region is conformationally heterogeneous and lacks stable receptor contacts, consistent with previous findings that the C-terminal loop is dispensable for AgRP binding^47^.

### A shared mechanism for endogenous inverse agonism at melanocortin receptors

Among the five melanocortin receptor subtypes, MC4R, like MC1R, exhibits constitutive Gs signaling activity sustained by an N-terminal tethered agonist^9,15^. AgRP suppresses this basal activity through a mechanism analogous to that of ASIP at MC1R. Similar to ASIP, the AgRP cysteine-rich domain functions as a molecular cork over the MC4R extracellular vestibule, sterically occluding access by both melanocortin peptides and the receptor’s N-terminus (Fig. 4a).

**Fig. 4.**
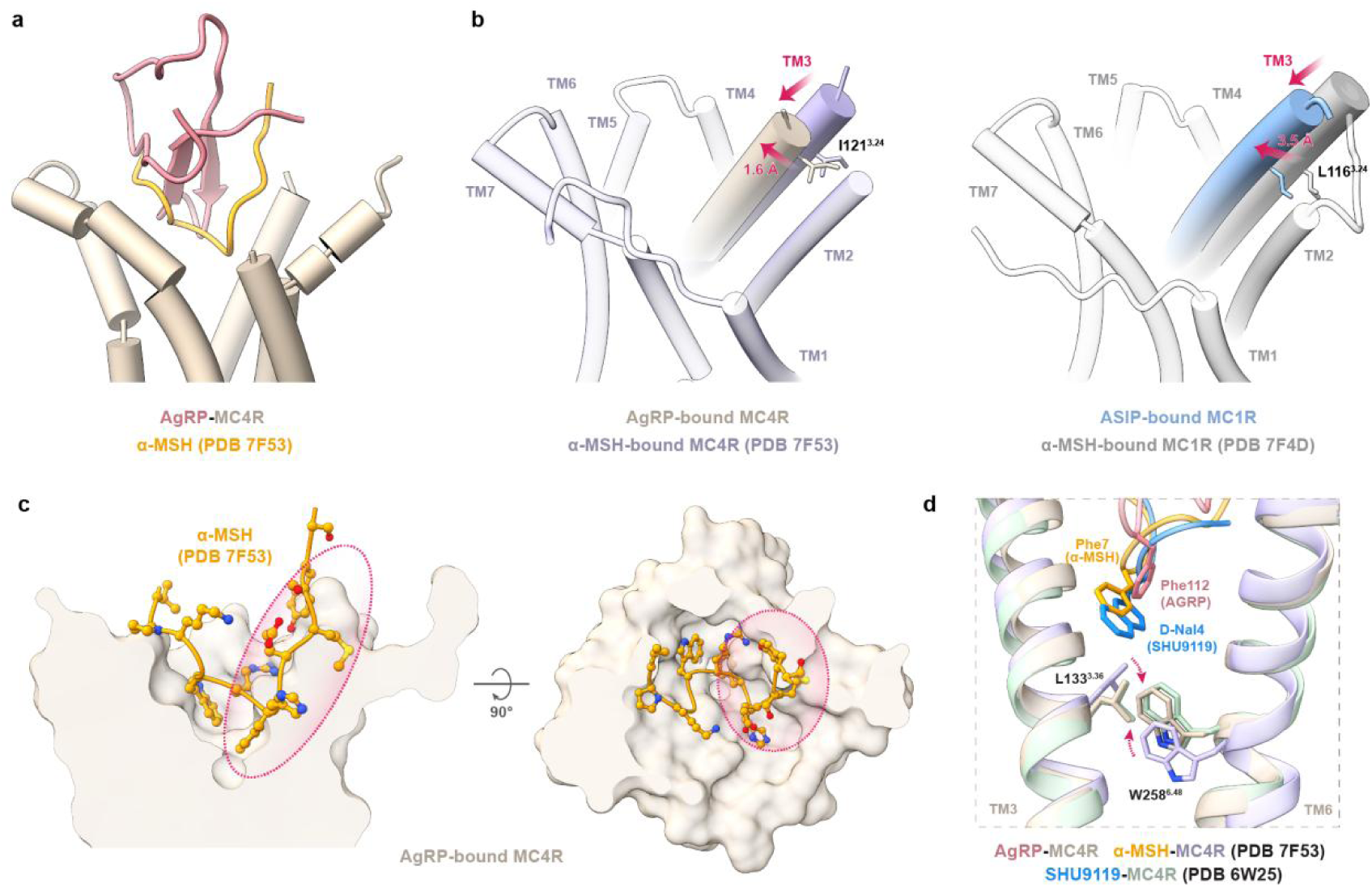
A conserved mechanism of inverse agonism by AgRP at MC4R. **a,** Superposition of the AgRP (red) and α-MSH (orange; PDB 7F53) binding positions within the orthosteric pocket, shown relative to the AgRP-bound MC4R (tan). **b,** Inward displacement of the extracellular end of TM3 in the inverse-agonist-bound states relative to the α-MSH-bound states. Left, AgRP-bound MC4R versus α-MSH-bound MC4R (purple), with TM3 shifting inward by 1.6 Å at I121^3.24^. Right, ASIP-bound MC1R (blue) versus α-MSH-bound MC1R (gray, PDB 7F4D), with TM3 shifting inward by 3.5 Å at L116^3.24^. Arrows indicate the direction of displacement. **c,** α-MSH superimposed onto the surface of the AgRP-bound MC4R orthosteric pocket, shown in two orthogonal views. The dashed ellipse marks the steric clash between α-MSH and the AgRP-constricted pocket. **d,** Comparison of the toggle-switch across the AgRP-MC4R, α-MSH-MC4R and SHU9119-MC4R (green, PDB 6W25) complexes. Phe112 of AgRP, Phe7 of α-MSH and D-Nal4 of SHU9119 are shown as sticks, together with the toggle residues L133^3.36^ and W258^6.48^.

Beyond this steric occlusion, AgRP binding is accompanied by remodeling of the extracellular receptor architecture (Extended Data Fig. 11a), recapitulating the hallmark rearrangement observed in ASIP-bound MC1R. In particular, TM3 shifts inward by 1.6 Å at I121^3.24^ relative to the α-MSH-bound state, in the same direction as the corresponding displacement in MC1R (3.5 Å at L116^3.24^) (Fig. 4b). By contrast, the antagonist SHU9119 displaces TM3 outward at the same position (Extended Data Fig. 11b). Despite the smaller displacement, superposition with α-MSH-bound MC4R confirms that even this reduced TM3 inward movement generates steric clashes with Tyr2, Met4 and Arg8 of α-MSH, constricting the agonist-contact face of the orthosteric pocket (Fig. 4c).

In addition to the conserved TM3 inward displacement, AgRP shares with ASIP a shallow mode of orthosteric engagement in which the RFF pharmacophore does not contact the toggle-gating residue L133^3.36^ (Fig. 4d). As in MC1R, the inward shift of extracellular TM3 is propagated along the helix, translating the L133^3.36^ Cα downward by 1.4 Å. Consequently, W258^6.48^ remains in its inactive orientation, and downstream microswitches adopt their inactive conformations (Fig. 4d and Extended Data Fig. 11c). This TM3-relayed mechanism contrasts with that of the synthetic MC4R antagonist SHU9119, whose D-Nal4 side chain penetrates substantially deeper into the receptor core to directly reposition L133^3.36^ and block the rotameric transition of W258^6.48^ (Fig. 4d). Thus, rather than inhibiting receptor activation through direct microswitch interference^48-51^ (Extended Data Fig. 11d), both AgRP and ASIP suppress constitutive Gs signaling through a multilayered mechanism integrating extracellular vestibule occlusion, arginine-for-calcium substitution, TM3-mediated pocket constriction, and helix-relayed toggle repositioning.

### Peripheral structural determinants of receptor subtype selectivity among melanocortin inverse agonists

Although both ASIP and AgRP function as endogenous melanocortin inverse agonists, they exhibit distinct receptor preference profiles^1,2^. AgRP potently inhibits both MC3R and MC4R but only weakly and incompletely inhibits MC1R (Fig. 5b), whereas ASIP primarily targets MC1R yet retains appreciable inhibitory activity at MC4R (Fig. 5a). Comparison of the ASIP-MC1R and AgRP-MC4R complexes reveals that the RFF pharmacophore is recognized through a highly conserved orthosteric recognition mode in both systems (Extended Data Fig. 9c,d), indicating that subtype preference is unlikely to arise from the orthosteric pharmacophore alone, but instead reflects differences in receptor tolerance. MC4R accommodates both ligands, whereas productive inhibition at MC1R is achieved only by ASIP. The key question is which peripheral structural features allow ASIP, but not AgRP, to meet the specific requirements of MC1R.

**Fig. 5.**
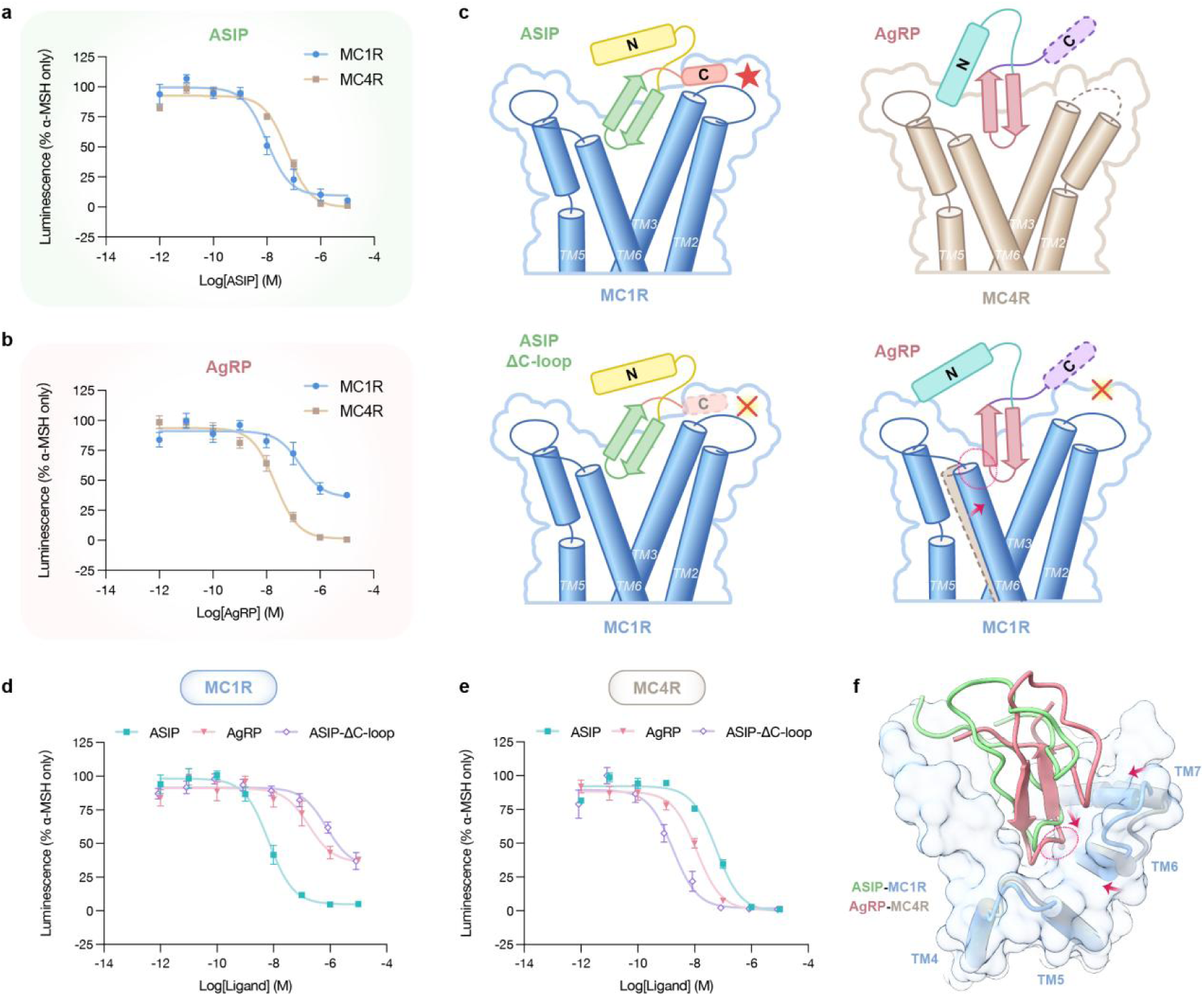
Peripheral structural determinants of ASIP and AgRP receptor selectivity. **a,** Dose-response curves of ASIP-mediated inhibition of α-MSH-stimulated signaling at MC1R (blue) and MC4R (tan), measured by GloSensor cAMP assay. **b,** Dose-response curves of AgRP-mediated inhibition of α-MSH-stimulated signaling at MC1R and MC4R, measured as in (**a**). **c,** Schematic model of C-terminal loop-dependent receptor selectivity of agouti proteins. At MC1R (blue), the ordered ASIP C-terminal loop (C) forms a clasp against TM2 and ECL1 that supports inhibition (top left), whereas its deletion (ASIP-ΔC-loop) abolishes this engagement (bottom left). At MC4R (tan), AgRP inhibits the receptor without an ordered C-terminal loop (top right). At MC1R, the disordered AgRP C-terminal loop cannot form the equivalent clasp, and the AgRP-bound pose is further incompatible with the intrinsically constricted TM6 of MC1R (bottom right). N, N-terminal loop; C, C-terminal loop. **d,e,** Dose-response curves of ASIP, AgRP, and ASIP-ΔC-loop inhibition of α-MSH-stimulated signaling at MC1R (**d**) and MC4R (**e**), measured as in (**a**). **f,** Superposition of the ASIP-MC1R and AgRP-MC4R complexes viewed from the extracellular side, showing the AgRP central loop (red) shifted toward the TM5/TM6 face relative to ASIP (green) with the MC1R TM6 shifted inward. Arrows indicate the displacement. The dashed ellipse marks the region of steric incompatibility with the constricted MC1R TM6. Data are shown as mean ± SEM from at least 3 biologically independent experiments for (**a**), (**b**), (**d**) and (**e**).

Structural comparison identifies the C-terminal loop as the principal structural determinant underlying the distinct receptor selectivity of ASIP and AgRP. In the ASIP-MC1R complex, this loop forms an ordered clasp against the extracellular tip of TM2 and ECL1 (Fig. 1j, Fig. 5c, top left and Extended Data Fig. 3g), whereas the extended C-terminal loop of AgRP remains disordered in the MC4R-bound structure and makes no defined receptor contacts (Fig. 3j and Fig. 5c, top right). Deleting this loop (ASIP-ΔC-loop) was sufficient to switch the receptor preference of ASIP (Fig. 5c, bottom left, d,e). At MC1R, the deletion reduced inhibitory potency by ∼100-fold and rendered inhibition incomplete, leaving ∼30% residual signaling (Fig. 5d). The resulting pharmacological profile closely overlapped that of AgRP at MC1R, effectively collapsing the functional distinction between the two ligands at this receptor. By contrast, the same deletion preserved, and even modestly enhanced, inhibition at MC4R (Fig. 5e). These findings identify the ASIP C-terminal loop as the defining structural element that confers full inhibitory efficacy at MC1R while remaining dispensable for inhibition of MC4R. The enhancement of activity at MC4R further argues that the loss at MC1R reflects a specific mechanistic requirement rather than compromised folding of the truncated protein.

This functional switch is explained by distinct modes of C-terminal loop engagement. In ASIP-bound MC1R, the ordered C-terminal clasp stabilizes an outward displacement of the TM2-ECL1 region, which is coupled to the pronounced inward movement of TM3 (Fig. 1j and Fig. 2c). By contrast, no equivalent C-terminal loop-mediated interaction is observed in the AgRP-MC4R complex (Fig. 3j and Extended Data Fig. 11a), and the associated inward displacement of TM3 is correspondingly attenuated (Fig. 4b). Together, these observations suggest that MC1R requires an additional C-terminal loop-mediated conformational coupling that reinforces the central loop-driven inward displacement of TM3 to stabilize the fully inhibited conformation. ASIP fulfills this requirement through its ordered C-terminal clasp, whereas AgRP lacks this additional interaction and therefore inhibits MC1R only weakly and incompletely, consistent with earlier loop-swap studies^28^. Accordingly, removing the C-terminal loop alone is sufficient to convert ASIP into an AgRP-like inhibitor of MC1R (Fig. 5d), establishing this single peripheral loop as sufficient to encode the receptor selectivity of ASIP.

Beyond the absence of C-terminal loop engagement, the AgRP-bound conformation itself is poorly compatible with the extracellular architecture of MC1R (Fig. 5c, bottom right). In the AgRP-MC4R complex, the central loop of AgRP adopts a position shifted toward the TM5-TM6 face of the receptor relative to the ASIP central loop in the ASIP-MC1R complex (Fig. 3b). However, the extracellular end of TM6 is positioned 2.9 Å further inward in MC1R than in MC4R (at the I/Y^6.58^ Cα), which would bring the AgRP central loop into direct steric conflict with the constricted TM6 face (Fig. 5f and Extended Data Fig. 11e). This constriction is already present in the ligand-free Nb18-stabilized state (3.3 Å at the same position) and is therefore intrinsic to the receptor rather than induced by ASIP (Extended Data Fig. 11e). Accommodating the TM5/TM6-oriented AgRP conformation within MC1R would therefore require additional helical remodeling, raisin<u>g</u> the structural barrier to productive engagement.

Together, these observations establish the divergent extracellular receptor surface, rather than the conserved orthosteric pocket, as the primary determinant of subtype selectivity among melanocortin inverse agonists (Fig. 5c). This selectivity is inherently asymmetric: MC1R constitutes a structurally restrictive receptor that imposes both a requirement for C-terminal loop-mediated coupling and an intrinsic TM6 constraint, whereas MC4R provides a comparatively permissive extracellular architecture that accommodates both inverse agonists.

### Identification of Nb96 as a selective neutral antagonist of MC1R

Although the agouti proteins are potent and selective endogenous inhibitors, their multiple disulfide bonds have historically complicated the production of homogeneous, correctly folded material^21^. This has motivated the search for scaffolds that preserve subtype selectivity while offering greater tractability. Single-domain nanobodies offer a compelling alternative, with favorable properties in production, stability, and tissue penetration^52^. Within the melanocortin family, this potential has so far been realized only for MC4R, for which a selective nanobody was recently described^40^, whereas no comparable selective modulator has been reported for MC1R. We accordingly screened for inhibitory nanobodies targeting MC1R.

An initial selection campaign using a fully synthetic yeast-display nanobody library^42^ yielded only non-inhibitory binders, exemplified by Nb18 (Extended Data Fig. 5a,b). We therefore returned to the immune library from which Nb1 had previously been isolated (Extended Data Fig. 1a). To enrich binders recognizing the extracellular pocket of the receptor, we implemented a dual-antigen cross-screening strategy during the yeast surface display selection (Fig. 6a). We alternated the selection rounds between two engineered MC1R constructs that share an identical extracellular region but have different intracellular fusion partners. One was the MC1R/KOR-Nb6 fusion used to identify Nb1, and the other was a de novo designed MC1R-Clip fusion construct^53^ (Extended Data Fig. 12a). By alternating between these two constructs, we specifically enriched binders recognizing the shared extracellular surface (Extended Data Fig. 12b). The final enriched population converged strongly onto a single dominant clone, designated Nb96, accompanied by a cluster of closely related variants differing primarily within CDR1 and CDR3 (Extended Data Fig. 12c).

**Fig. 6.**
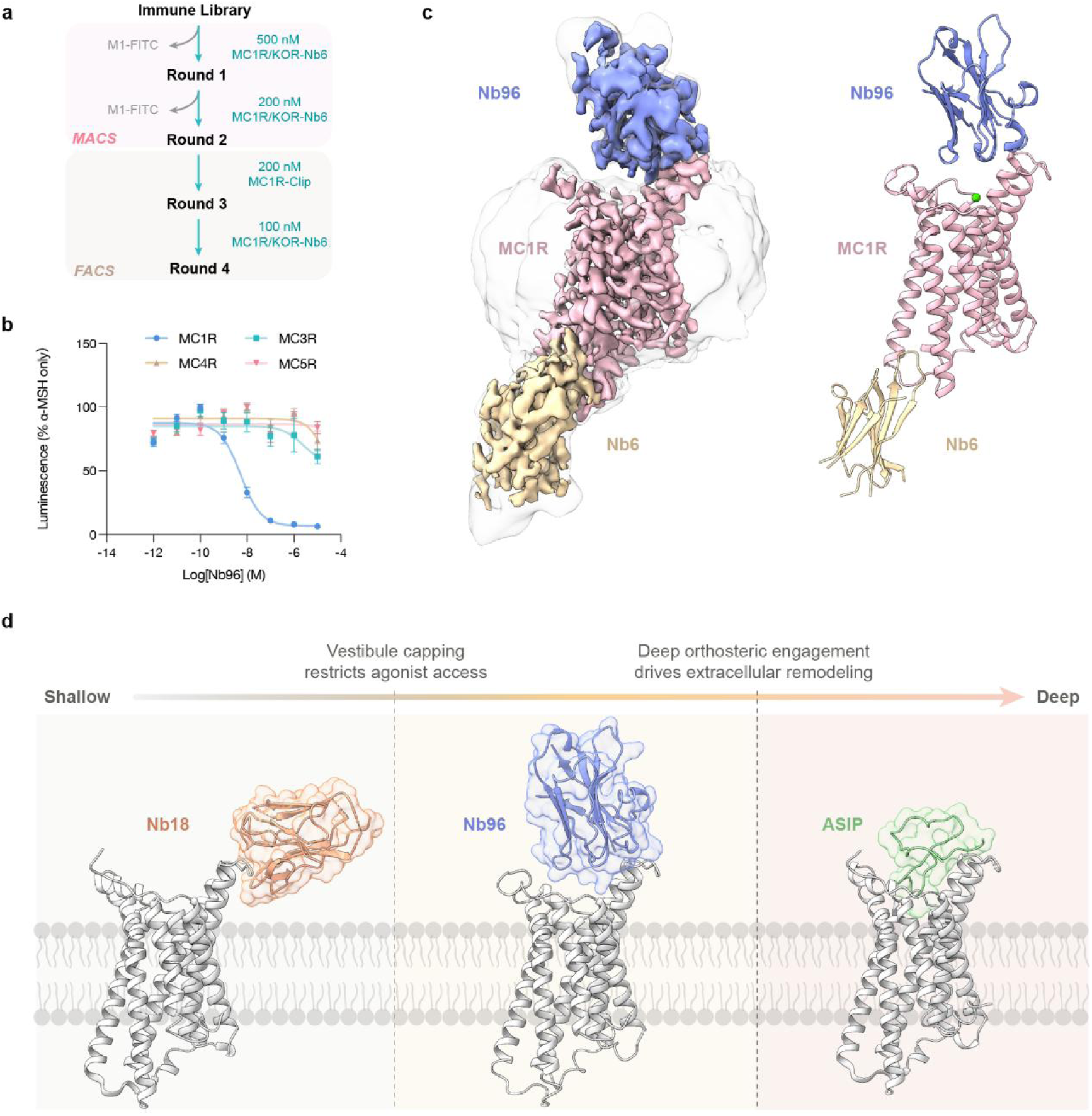
Nb96 is an MC1R-selective neutral antagonist that caps the extracellular vestibule. **a,** Dual-antigen cross-screening schematic for isolation of extracellular surface targeting nanobodies. **b,** Inhibition of α-MSH-stimulated signaling by Nb96 at MC1R, MC3R, MC4R and MC5R, measured by GloSensor cAMP assay and normalized to the α-MSH-only response. Data are shown as mean ± SEM from 6 biologically independent experiments for MC1R and 3 biologically independent experiments for MC3R, MC4R and MC5R. **c,** Cryo-EM density map (left) and model (right) of the Nb96-MC1R complex. Nb96, orchid; MC1R, pink; Nb6, yellow. The detergent micelle is shown as a transparent surface and the orthosteric Ca^2+^ as a green sphere. **d,** Engagement depth comparison of three extracellular MC1R ligands, arranged from shallow to deep: Nb18 (functionally silent; peripheral ECL1 engagement), Nb96 (neutral antagonist; vestibule capping without orthosteric penetration) and ASIP (inverse agonist; deep insertion).

Nb96 bound MC1R with an apparent affinity of ∼7.5 nM measured by on-yeast titration (Extended Data Fig. 12e) and inhibited α-MSH-stimulated MC1R cAMP signaling with an IC_50_ of ∼5.1 nM in cell-based GloSensor assays (Fig. 6b). Nb96 showed no detectable effect on the receptor constitutive activity (Extended Data Fig. 12f), identifying it as a neutral antagonist that inhibits agonist-stimulated MC1R signaling. Nb96 also displayed marked subtype selectivity, with markedly reduced potency at MC3R and no measurable effect at MC4R or MC5R (Fig. 6b). Closely related variants of Nb96 exhibited comparable inhibitory potency at MC1R, with IC_50_ values clustered around 10 nM (Extended Data Fig. 12d), consistent with functional convergence onto a shared binding mode. Nb96 was therefore selected as the representative clone for subsequent structural and mechanistic characterization.

### Structural basis of Nb96-mediated neutral antagonism at MC1R

To define the molecular mechanism of Nb96-mediated inhibition, we determined the cryo-EM structure of the Nb96-MC1R complex at a global resolution of 3.07 Å (Fig. 6b, Extended Data Fig. 12g, Extended Data Fig. 13 and Table 1). The Nb96-bound MC1R adopts a canonical inactive conformation, with TM6 in its inward position and the transmembrane bundle closely matching the Nb18-bound ligand-free state (Cα RMSD < 0.8 Å; Extended Data Fig. 14a). Rather than inserting a CDR loop into the orthosteric pocket, Nb96 engages the receptor surface adjacent to ECL1 and sits as a shallow cap over the extracellular vestibule (Fig. 6c). This contrasts with most extracellular-facing class A GPCR nanobodies, whose CDR3 loops typically penetrate deeply into the orthosteric pocket to engage and modulate the receptor^37–40,54–59^ (Extended Data Fig. 15). Superposition with the α-MSH-bound state reveals a steric overlap between the Nb96 CDR1 loop and the N-terminal segment of α-MSH above the ligand-binding pocket (Extended Data Fig. 14b). This peripheral, vestibule-capping mode accounts for the ability of Nb96 to block agonist-stimulated signaling while leaving constitutive receptor activity intact.

Comparing Nb96 to the functionally silent Nb18 reveals how binding depth affects receptor activity. Nb96 caps the top of the orthosteric binding pocket, whereas Nb18 only binds to the peripheral edge of ECL1 with most of its body exposed laterally to solvent (Fig. 6d). Consistently, Nb18 exhibits no detectable effect on MC1R signaling (Extended Data Fig. 5b). Nb96 also differs from ASIP, which inserts deeply into the binding pocket (Fig. 6d). Comparing Nb18, Nb96, and ASIP shows a clear progression: peripheral ECL1 binding is functionally silent (Nb18), capping the vestibule without orthosteric penetration causes neutral antagonism (Nb96), and deep insertion into the pocket drives inverse agonism (ASIP). These observations reveal that simply blocking the vestibule is insufficient to suppress MC1R activity; deeper insertion into the orthosteric pocket is required to suppress constitutive signaling (Fig. 6d).

### Molecular determinants of MC1R-selective recognition by Nb96

The Nb96-MC1R interface involves all three CDR loops and centers on extensive contacts with ECL1 and the extracellular tips of TM2 and TM3 (Fig. 7a). At the core of this interface, CDR3 forms an antiparallel β-strand against ECL1 (Fig. 7b). This geometric framework is anchored by two backbone hydrogen bonds between Tyr102 of CDR3 and A108^ECL1^, and between Asp100 of CDR3 and A110^3.18^ (Fig. 7b). Two polar interaction nodes positioned at opposite ends stabilize this β-sheet scaffold. At the N-terminal edge of ECL1, E102^2.68^ forms hydrogen bonds with CDR3 (Tyr102 and Gly105) and CDR2 (Ser54) (Fig. 7c). At the opposite end, Q114^3.22^ is clamped by Ser28 of CDR1 through both backbone and side-chain hydrogen bonds (Fig. 7d). Interestingly, both E102^2.68^ and Q114^3.22^ also interact with ASIP, though in different ways (Extended Data Fig. 14c). ASIP packs hydrophobically against E102^2.68^ via Val127 and clamps Q114^3.22^ using bidentate hydrogen bonds with the Cys116 backbone. The convergent recruitment of E102^2.68^ and Q114^3.22^ by two structurally unrelated ligands identifies these residues as central interaction hubs within the MC1R vestibule. Surrounding these polar nodes, hydrophobic interactions further strengthen the interface (Fig. 7e). Trp101 of CDR3 packs against V107^ECL1^, while Tyr31 and Tyr32 of CDR1, together with Tyr102 of CDR3, form extensive aromatic and hydrophobic interactions with I98^2.64^, L101^2.67^, A110^3.18^, and L113^3.21^. Alanine substitutions at key interfacial residues on Nb96 or MC1R reduced Nb96 inhibitory potency in cAMP assays (Fig. 7f,g; Extended Data Fig. 12h), confirming the functional importance of this network.

**Fig. 7.**
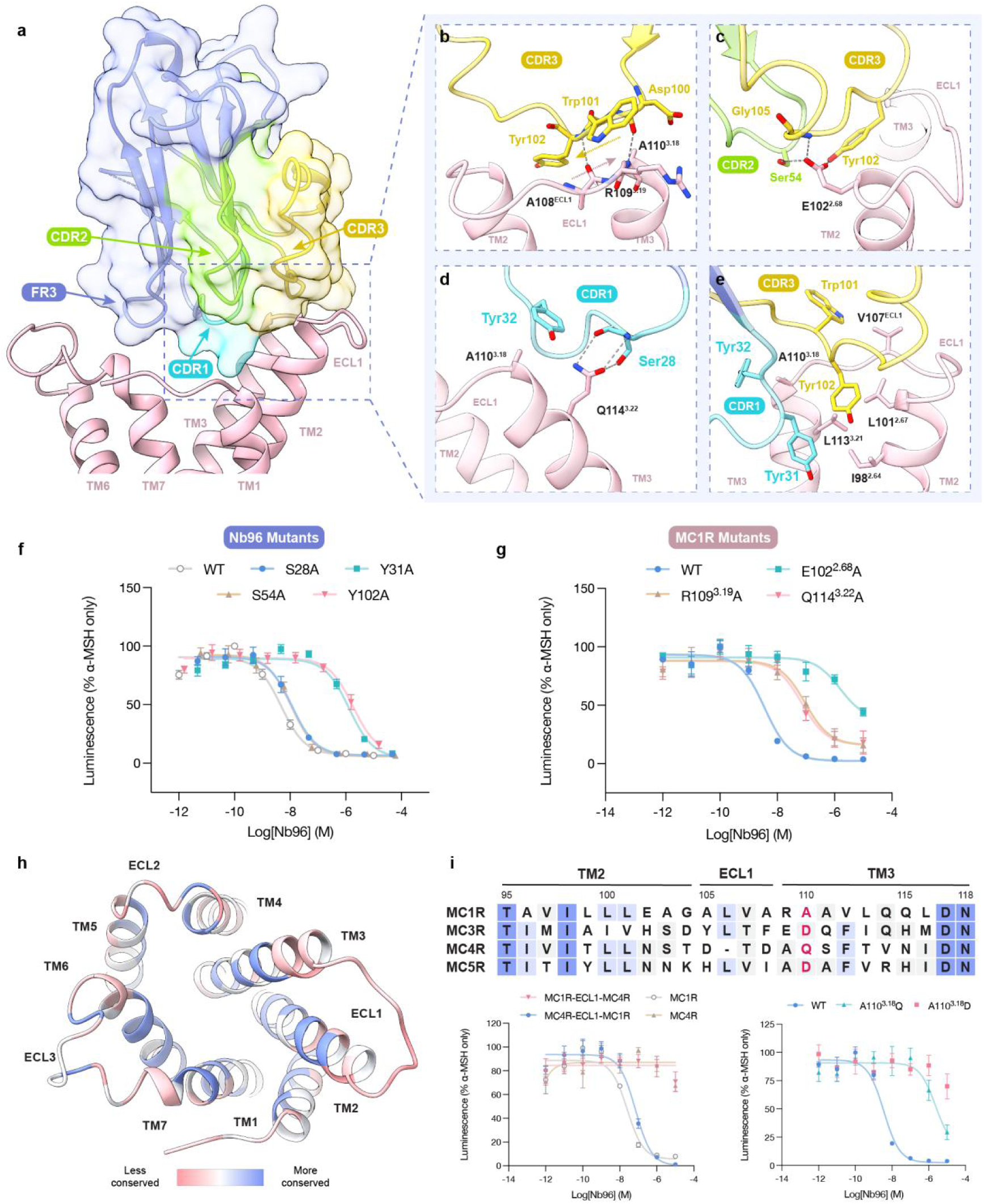
An ECL1-centered interface underlies selective recognition of MC1R by Nb96. **a,** Overall view of the Nb96-MC1R interface, formed by the three CDR loops of Nb96. CDR1, cyan; CDR2, green; CDR3, yellow; FR3, orchid; MC1R, pink. **b-e,** Detailed views of the interaction nodes within the Nb96-MC1R interface. CDR3 engages ECL1 as an antiparallel β-strand (**b**). E102^2.68^ is engaged by Tyr102 and Gly105 of CDR3 and Ser54 of CDR2 (**c**). Q114^3.22^ is clamped by Ser28 of CDR1 (**d**). Hydrophobic packing of CDR1 (Tyr31, Tyr32) and CDR3 (Trp101, Tyr102) against ECL1, TM2 and TM3 (**e**). Interacting residues are shown as sticks and gray dashed lines indicate polar interactions. **f,g,** Effect of alanine substitutions in Nb96 (**f**) and MC1R (**g**) on Nb96 inhibitory potency. **h,** MC1R surface colored according to sequence conservation across the melanocortin receptor family (blue, more conserved; red, less conserved). **i,** Top, sequence alignment of MC1R, MC3R, MC4R and MC5R across TM2, ECL1 and TM3, with A110^3.18^ highlighted in red. Bottom left, Nb96 inhibition of MC1R, MC4R and the reciprocal ECL1-swap chimeras. Bottom right, Nb96 inhibition of MC1R and the A110^3.18^D and A110^3.18^Q mutants. Data in (**f**), (**g**) and (**i**) were measured by GloSensor cAMP assay, normalized to the α-MSH-only response, and are shown as mean ± SEM from at least 3 biologically independent experiments.

The ECL1-centered footprint of Nb96 further explains its subtype selectivity. Sequence alignment across the MCR family shows that while the orthosteric pocket residues are highly conserved, the extracellular surface residues surrounding ECL1 vary significantly (Fig. 7h,i, top). Consistent with this, Nb96 selectively inhibited MC1R yet displayed markedly weaker activity at MC3R and no effect at MC4R or MC5R (Fig. 6b). To confirm that this divergent region drives selectivity, we swapped the ECL1 segment of MC1R with the corresponding region from MC4R. This chimeric receptor was no longer inhibited by Nb96 (Fig. 7i, bottom left). Conversely, grafting the MC1R ECL1 segment onto MC4R made the previously insensitive receptor susceptible to Nb96 (Fig. 7i, bottom left). Together, these results establish that the ECL1-centered interface is both necessary and sufficient for subtype-selective recognition. We next identified A110^3.18^, a residue at the extracellular edge of TM3, as a key factor. In the Nb96 binding site, A110^3.18^ helps form both the intermolecular β-sheet scaffold and the surrounding hydrophobic packing (Fig. 7b,d,e). This position is uniquely occupied by alanine in MC1R, whereas the subtypes carry either aspartate or glutamine (Fig. 7i, top). Substitution of A110^3.18^ with either aspartate or glutamine abolished or severely reduced Nb96 inhibitory activity (Fig. 7i, bottom right), confirming this position as a critical selectivity hotspot within the ECL1 interface.

## Discussion

Constitutive activity is a widespread feature of GPCR signaling^60^, yet endogenous inverse agonists remain rare. This suggests that suppressing baseline signaling is only necessary under specific physiological conditions. The melanocortin system is unique in containing two independent endogenous inverse agonists. In pigmentation, ASIP drives down the constitutive activity of MC1R to enable the switch from eumelanin to pheomelanin^4,33^, whereas in energy homeostasis, AgRP suppresses the constitutive signaling of MC4R to promote feeding^5,34^. The independent emergence of endogenous inverse agonism in these two melanocortin systems underscores a physiological requirement for active suppression below the constitutive signaling state that could not be achieved through receptor desensitization^61^. However, whether endogenous inverse agonists converge on a common structural mechanism for receptor silencing, and how receptor selectivity is simultaneously achieved, have remained unresolved. By determining the first structures, to our knowledge, of GPCRs bound to endogenous inverse agonists, we establish the structural principles of this specialized mode of GPCR regulation, revealing both the shared mechanism and the divergent features of endogenous melanocortin inhibition.

Despite targeting different melanocortin receptors, ASIP and AgRP employ a conserved structural strategy for receptor recognition and inverse agonism. In both complexes, the cysteine-rich domain of each agouti protein caps the extracellular vestibule as a molecular cork, while its RFF pharmacophore fits in the orthosteric pocket without inducing receptor activation. An arginine side chain replaces the activating calcium ion within the conserved acidic triad, anchoring a shallow insertion without triggering the conserved microswitch rearrangements associated with receptor activation. Receptor silencing is imposed from the extracellular surface, where the cysteine-rich domain remodels the extracellular structure, driving an inward displacement of TM3 that constricts the orthosteric pocket and propagates along the helix to hold the toggle-gating residue inactive without direct contact. This mechanism is different from that employed by the reported small-molecule antagonists, in which ligands typically insert deeply into the transmembrane core to directly engage the conserved toggle switch, thereby restraining the outward movement of TM6 associated with GPCR activation^46^. The conservation of this indirect, TM3-relayed mechanism across both receptors implies that subtype selectivity of the endogenous inverse agonists is achieved not by the silencing mechanism itself, but by targeting the divergent extracellular surface.

Our structures and functional data identify the ASIP C-terminal loop as the principal determinant of subtype selectivity. Unlike the disordered C-terminal loop of AgRP, the ordered ASIP loop forms an extensive interface with the extracellular tip of TM2 and ECL1, and its deletion renders ASIP an AgRP-like, partial inhibitor of MC1R. Whereas the central loop of AgRP is sufficient to silence MC4R, complete silencing of MC1R appears to additionally depend on the ASIP C-terminal loop as a structural buttress that enables the larger inward displacement of TM3. This differential requirement may originate in intrinsic properties of the receptors. MC1R displays the highest constitutive activity among melanocortin receptors^8^, such that its complete silencing likely requires a larger extracellular conformational rearrangement. Notably, this seems to depend less on individual key contacts and more on how well the ordered loop complements the overall shape of the receptor surface. Therefore, subtype selectivity is mainly determined by the divergent extracellular surface rather than the conserved orthosteric pocket. This separation provides a general framework for achieving subtype selectivity toward GPCR subtypes sharing highly conserved orthosteric pockets.

The Nb18 and Nb96 structures further establish the extracellular receptor surface as a key determinant of both pharmacological efficacy and subtype selectivity. Together with ASIP, these binders demonstrate that distinct modes of engagement can produce a continuum of pharmacological outcomes, ranging from functional silence to neutral antagonism and inverse agonism. Notably, despite their distinct modes of engagement and functional effects, all three ligands engage the same ECL1-centered vestibule. The convergence of endogenous ligands and engineered nanobodies on this region highlights it as a tunable regulatory interface. Consequently, targeting this extracellular interface represents a promising strategy for designing modulators that simultaneously tune efficacy and selectivity, a longstanding challenge for receptors with highly conserved orthosteric pockets.

